# CD300f is required for efficient lipid handling during cuprizone-induced demyelination

**DOI:** 10.64898/2026.11.05.764483

**Authors:** Gemma Manich, Natalia Rego, Elvira de Frutos, Paula Sanchez-Molina, Beatriz Almolda, Rubèn López-Vales, Berta González, Bernardo Castellano, Hugo Peluffo, Natalia Lago

**Author notes:** **Corresponding authors:** Dr. Natalia Lago, Department of Cell Biology, Physiology and Immunology, Institut de Neurociències, Universitat Autònoma de Barcelona, 08193 Bellaterra, Barcelona, Spain, Dr. Gemma Manich, Department of Morphological Sciences. Facultat de Medicina, Institut de Neurociències, Universitat Autònoma de Barcelona, 08193 Bellaterra, Barcelona, Spain.

## Abstract

Microglia play a crucial role in central nervous system homeostasis and response to demyelination, yet the mechanisms that coordinate the sensing and processing of myelin-derived lipids remain poorly understood. CD300f is a lipid-sensing innate immune receptor expressed by myeloid cells that has been implicated in tissue damage sensing, phagocytosis, and lipid metabolism. Here, we investigated the role of CD300f in the cuprizone-induced demyelination model. Using CD300f-deficient mice, we demonstrate exacerbated demyelination, increased lipid droplet accumulation, impaired microglial responses, and reduced remyelination accompanied by motor coordination deficits. Bulk RNA sequencing of the corpus callosum at the peak of demyelination revealed enrichment of homeostatic, disease-associated (DAM), and foamy microglial signatures, together with increased expression of genes involved in lysosomal function, phagocytosis, inflammation, and lipid metabolism. Notably, these transcriptional changes were associated with marked lipid droplet accumulation in microglia and enrichment of microglial states previously linked to chronic demyelinating lesions. These findings suggest that CD300f is required for efficient handling of myelin-derived lipids and for microglial metabolic adaptation during demyelination. Our study identifies CD300f as a critical regulator of microglial function and lipid homeostasis and highlights this receptor as a potential therapeutic target in demyelinating diseases.

## 1. INTRODUCTION

Microglia are the resident macrophages of the central nervous system (CNS) and exert diverse functions essential for brain development, adult homeostasis, and disease. These cells express a wide array of immune receptors that detect perturbations in the CNS milieu, such as cellular debris, pathogens, or blood–brain barrier disruption, and in response, they rapidly act to restore tissue integrity (Davalos et al., 2005; Nimmerjahn et al., 2005). Among the various perturbations, microglia respond robustly to demyelination, as occurs in multiple sclerosis (MS) pathology, a chronic autoimmune inflammatory disorder (Franklin and ffrench-Constant, 2017; Lloyd and Miron, 2019; Franklin and Simons, 2022). In this context, microglia undergo marked phenotypic changes to adapt to the altered environment and acquire new functions aimed at restoring homeostasis (Keren-Shaul et al., 2017; Hammond et al., 2019).

In MS, remyelination constitutes an endogenous regenerative response to progressive demyelination; however, the mechanisms underlying remyelination failure or inefficiency in MS remain incompletely understood. An increased prevalence of active and mixed active/inactive white matter lesions harboring foamy microglia has been associated with accelerated disease progression (van der Vliet et al., 2026). This relationship was absent in patients with lesions containing non-foamy microglia, while a greater abundance of remyelinated lesions correlated with slower progression, suggesting an important link between active demyelination, microglia/macrophages lipid metabolism and functional compromise. Microglia can phagocytose myelin debris generated during demyelination via several receptors, including CD36, TAM receptors (Tyro3, Axl, and Mer), Fc receptors, and the lipid-binding immune receptor TREM2 (Gao et al., 2024). TREM2 has been shown to regulate microglial responses to demyelination in the cuprizone (CPZ) mouse model and in bone marrow–derived macrophage (BMDM) cultures (Damisah et al., 2020; Nugent et al., 2020). Furthermore, TREM2 modulates the remyelination process following lysolecithin-induced demyelination (Wang et al., 2023), and enhancing TREM2 function improves lipid clearance by phagocytes in spinal cord demyelination (Bosch-Queralt et al., 2021).

Another receptor putatively involved in lipid phagocytosis and processing is the lipid-binding immune receptor CD300f. CD300f belongs to the CD300 family, which comprises eight members classified as either activating or inhibitory immune receptors (Borrego, 2013). Uniquely, CD300f can deliver activating signals through motifs that bind the p85α regulatory subunit of PI3K and growth factor receptor–bound protein 2 (Grb2). This activating signaling capacity is critical for phosphatidylserine-mediated macrophage efferocytosis (Tian et al., 2014).

Under physiological conditions, CD300f is expressed by microglia and other myeloid cells and functions as a dual activating/inhibitory receptor involved in the phagocytosis of apoptotic cells and potentially synapses (Tian et al., 2014; Lago et al., 2020; Negro-demontel et al., 2024). Notably, CD300f shares several ligands with TREM2, including phospholipids such as phosphatidylserine, sphingomyelin and phosphatidylcholine, as well as ceramides and lipoproteins (Choi et al., 2011; Tian et al., 2014). Beyond ligand recognition, CD300f is emerging as an important regulator of myeloid cell metabolism. Its absence induces profound metabolic alterations in microglia, including reduced glycolysis, fatty acid oxidation, mitochondrial oxidative phosphorylation and autophagy under inflammatory conditions (Lago et al., 2020). Similar defects have been reported in CD300f-deficient bone marrow-derived macrophages (Evans et al., 2023), suggesting a broader requirement for CD300f in maintaining metabolic fitness and adaptive metabolic reprogramming. Consistently, loss of CD300f promotes the emergence of a damage-associated microglia (DAM) signature during aging (Evans et al., 2023), highlighting its role in regulating microglial functional states.

These properties position CD300f as a potential regulator of microglial responses to damage-associated molecular patterns (DAMPs) and lipid-rich debris. Consistent with this idea, a genome-wide screen aimed at identifying ITIM-containing receptors relevant to multiple sclerosis pathology identified CD300f as the most upregulated gene in experimental autoimmune encephalomyelitis (EAE). Moreover, genetic deletion of CD300f exacerbated neuroinflammation and worsened neurological deficits in this model (Xi et al., 2010). Similar protective effects have been reported in several peripheral inflammatory and autoimmune disorders, including allergy, systemic lupus erythematosus, colitis and LPS-induced skin inflammation (Izawa et al., 2012; Tian et al., 2014; Moshkovits et al., 2015, 2017). Together, these findings support a key role for CD300f in limiting inflammatory damage and maintaining myeloid cell homeostasis across tissues.

In the present study, we investigated the role of CD300f in demyelination and remyelination using the CPZ model of MS (Matsushima and Morell, 2001). This model is particularly well suited for assessing microglia-intrinsic mechanisms, as it does not involve infiltration of peripheral monocytes (Mildner et al., 2007). Using this approach, we demonstrate that CPZ-induced demyelination is exacerbated in CD300f-deficient mice, accompanied by increased lipid droplet accumulation and more severe motor impairments compared with wild-type (WT) controls. These alterations are associated with dysregulation of genes linked to microglial states, metabolism, and immune activation.

## 2. METHODS

### 2.1. Animals

Adult (8-12 weeks old) male and female CD300f^−/−^ (Genentech (Xi et al., 2010)) and WT mice on the C57BL/6 background were used in this study. Mice were maintained with food and water *ad libitum* in a 12 h light/dark cycle during the experiments. All the experimental work was approved by the Universitat Autònoma de Barcelona Animal Care Committee (UAB-CEEA71) according to Spanish legislation (Ley 7/2023, Ley 32/2007, and Real Decreto 53/2013, as amended) and European legislation (Directive 2010/63/EU on the protection of animals used for scientific purposes).

### 2.2. Cuprizone diet

Demyelination was induced with CPZ intoxication, as described in Petkovic et al. (Petković et al., 2016). Briefly, WT and CD300f^−/−^ mice were fed with 0.2% (w/w) CPZ (bis–cyclohexanone-oxaldihydrazone, Sigma-Aldrich) mixed into standard powdered ground chow for 5 weeks. Animals were fed with regular powdered chow three days before starting the CPZ treatment. The CPZ diet was administered freshly made daily. After 3 or 5 weeks of CPZ treatment, mice were processed for either histochemistry/immunohistochemistry analysis or transcriptomics analysis (Figure 1a). To study remyelination, one group of animals was maintained on a regular diet for an additional week, after completing the 5-week CPZ treatment (Figure 1a). A group fed with regular diet without CPZ was added to the study and used as experimental control group for all the histological analysis.

**Figure 1:**
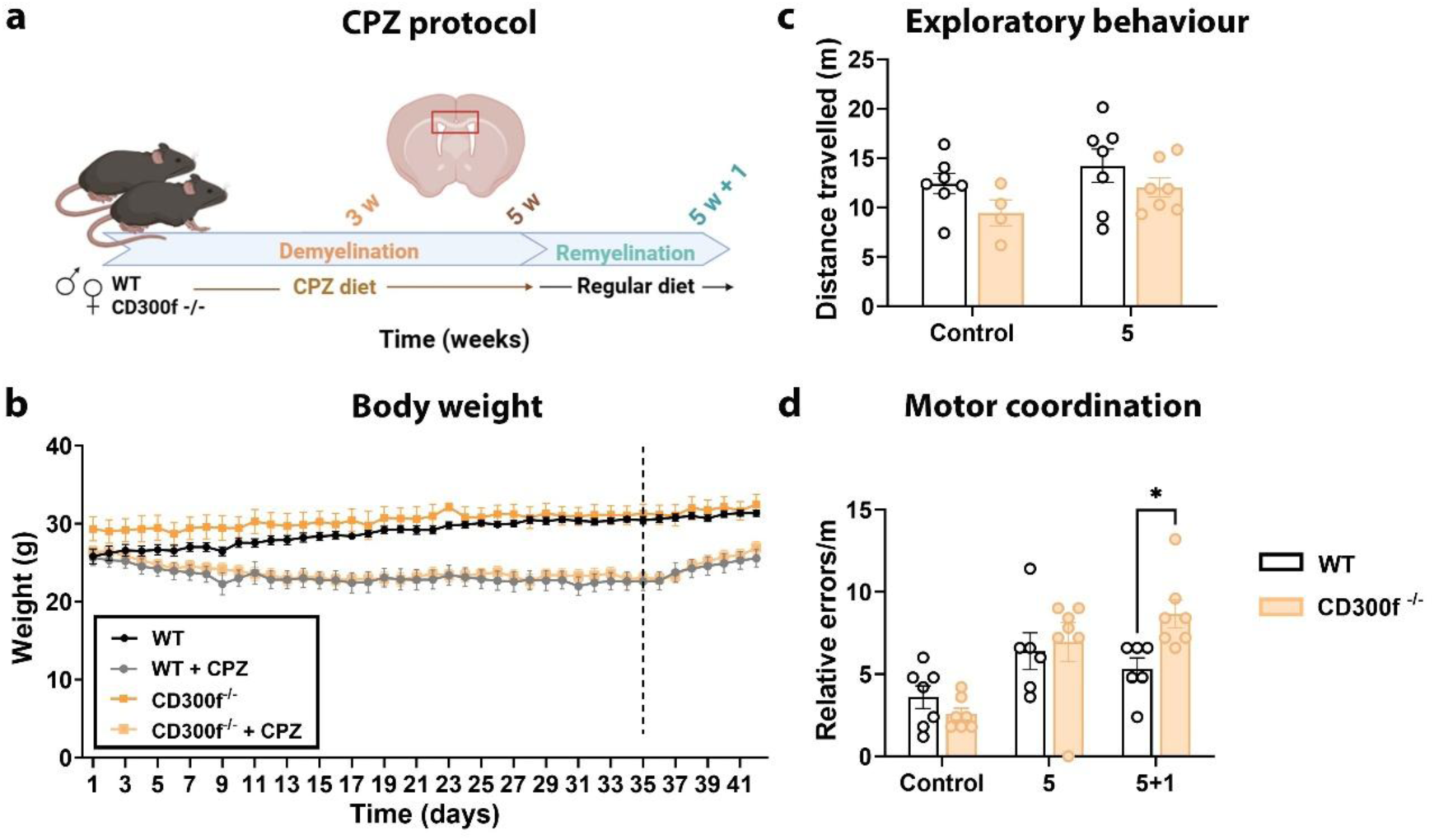
Absence of CD300f reduces motor coordination during remyelination. **(a)** Schematic representation of treatment paradigms in the cuprizone (CPZ) demyelination and remyelination model. **(b)** Graph of animal absolute body weight (g) in control and CPZ-treated WT and CD300f^−/−^ mice during the 5-week treatment of CPZ intoxication (day 1-35, demyelination), and after 1 week of CPZ withdrawal (day 36 to 42, remyelination). **(c)** Assessment of exploratory behavior by Open Field Test in control and 5-week CPZ-treated WT and CD300f**^−/−^** male mice. **(d)** Analysis of motor coordination by Grid test in control WT and CD300f**^−/−^** mice, after a 5-week CPZ-treatment (demyelination) and after 1 week of CPZ withdrawal (5+1, remyelination) in both genotypes. Data are presented as mean ± SEM. Each dot represents an animal. Statistical analysis: Two-way ANOVA plus Sidak post-hoc test; *p ≤ 0.05.

### 2.3. Behavioural analysis

Open Field Test (OFT). The OFT was performed to assess the locomotor activity and spontaneous activity of mice. The apparatus consisted of a white acrylic box measuring 40 cm in length × 60 cm in width × 50 cm in height. The animals were exposed for 6 min to the arena of the box and the total distance travelled was recorded and evaluated using the AnyMaze software. The arena was cleaned with 10% alcohol between animals in order to exclude possible clues.

Grid Test. Sensorimotor deficits following the cuprizone diet were assessed using a cage (40 cm in length × 27 cm in width × 30 cm in height) with a floor made of 1.6-mm diameter rods spaced 10 mm apart and elevated 1.2 cm above the base. Mice were placed on the cage grid for 1 minute (Grill et al., 1997). The time spent in movement and the number of incorrect paw placements were recorded. The number of incorrect paw placements was expressed relative to the time spent in movement. All video recordings were reviewed by three independent experimenters to ensure scoring accuracy.

### 2.4. Tissue processing for histological analysis

Animals were deeply anesthetized with ketamine (90 mg/kg, ip) and xylazine (10mg/kg, ip) and perfused intracardially for 10 min with 4% paraformaldehyde (PFA) in 0.1 M phosphate buffer (PB, pH 7.4). Brains were excised and immersed in the same fixative for either 24 h at 4°C, before dehydration and paraffin embedding, or for 4 h at 4°C, before being cryoprotected in 30% sucrose in 0.1 M PB for 48 h, frozen in isopentane (320404, Sigma-Aldrich) chilled in dry ice, and stored at -80°C. 10-µm-thick serial coronal brain sections from paraffin blocks were cut and collected on slides using a microtome Leitz 1512. Alternatively, 30-µm-thick free-floating serial coronal brain sections were cut with a cryostat (CM3050S Leica) and stored in an anti-freezing Olmos solution at -20°C until use.

### 2.5. Luxol Fast Blue staining

Myelin staining was performed as described by Petkovic et al. (Petković et al., 2016). Coronal brain paraffin sections were deparaffined, hydrated and incubated in Luxol Fast Blue solution (LFB; 0.1 g LFB/100 mL of 95% ethanol with 10.5 mL of 10% CH_3_COOH) for 3 h at 60°C. The excess was washed with 95% ethanol and distilled water, and the sections were differentiated using 0.05% Li_2_CO_3_ solution, followed by 70% ethanol. Sections were dehydrated in alcohol, cleared in xylene, and covered with a DPX mounting medium.

### 2.6. Oil Red O staining

Slides with brain cryostat sections were incubated in freshly made and filtered oil red O (ORO) solution (ORO stock solution: 2.5 g/400 mL 99% isopropyl alcohol; ORO working solution: 1.5 stock solution/1 distilled water) for 2 h at room temperature (RT) (Mehlem et al., 2013). Excess was washed with tap water, and the slides were cover-slipped with PermaFluor aqueous mounting medium (TA-030-FM, Thermo Fisher Scientific).

### 2.7. Immunohistochemistry on paraffin and cryostat sections

The immunohistochemical detection of Olig-2, GFAP, APC, Pu.1 and Iba-1 was performed in paraffin sections as follows: Sections were deparaffinized and after washing with distilled water, and when needed, were immersed in either a citrate buffer solution (pH=6.0) or (pH=8.0) at 98°C for 20 min for antigen retrieval (Supplementary Table 1). After washing with distilled water, the activity of endogenous peroxidase was blocked with a 5 min incubation of 2% H_2_O_2_ in 70% methanol and washed twice with 0.05 M Tris-buffered saline (TBS), pH 7.4 containing 0.1% Triton X-100 (TBS-0.1%T). Then, sections were incubated for 1 h in blocking buffer solution (BB; TBS-0.1%T containing 10% fetal calf serum and 3% bovine serum albumin (BSA)), followed by the incubation of primary antibodies diluted in BB (Supplementary Table 1) overnight at 4°C and 1 h at RT. After washing with TBS-0.1%T, sections were incubated for 1 h at RT with the corresponding secondary antibodies diluted in BB. Then, sections were washed with TBS-0.1%T and incubated in streptavidin-peroxidase for 1h (1:500; SA-5004; Vector Laboratories, Inc; Burlingame,CA). The reaction was visualized by incubating the sections with DAB kit (SK-4100; Vector Laboratories, Inc; Burlingame, CA) following the manufacturer’s instructions. All sections were air-dried, dehydrated in graded alcohols followed by xylene treatment, and coverslipped with DPX. Sections incubated in BB without the primary antibody were used as negative controls. Stained sections were observed with a Nikon Eclipse 80i bright-field microscope.

Immunohistochemical detection of dMBP was performed on free-floating sections (1:8000). The protocol was done as explained above, but using TBS-1%T buffer instead of TBS-0.1%T. After visualizing the reaction with DAB, sections were mounted, air-dried, and coverslipped with DPX. For double immunofluorescence staining, tissue sections were first incubated with the primary antibody against Iba1, followed by the appropriate fluorescent secondary antibody. Lipid droplets were subsequently labeled with BODIPY™ 493/503 (Thermo Fisher Scientific; 1:4000). Sections were counterstained with DAPI and mounted with antifade mounting medium before imaging by by a Leica TCS SP5 (Servei de Microscòpia i Raigs X, UAB) confocal fluorescence microscope.

### 2.8. Brightfield microscopy quantification and cell counting

Sections were observed and photographed using a DXM 1200F Nikon digital camera mounted on a Nikon Eclipse 80i bright-field microscope. For densitometric analyses, images from 2 to 4 sections containing the central corpus callosum (CC) were taken and analysed using the Analysis® software. For staining quantification, the percentage of the area occupied by staining was recorded. The density of Olig2+ cells, APC+ cells, and Pu.1+ cells (cells/mm^2^) was quantified manually using the “Cell counter” plug-in from NIH ImageJ software (Wayne Ras-band, National Institutes of Health, USA).

### 2.9. RNA sequencing

After 5 weeks of CPZ diet, 5 WT and 4 CD300f^−/−^ male mice were deeply anaesthetized with Pentobarbitone (200 mg/kg; ip) and perfused with cold PBS. CC was dissected out and homogenized in Primozol (SIGMA, T9424), and the resulting aqueous phase was purified using the Qiagen RNA MiniPrep Plus kit (Qiagen). After extraction, RNA quality was determined using the NanoDrop ND-1000 device (Thermo Fisher). Total RNA samples were processed at BGI Tech Solutions (Hong Kong). RNA integrity and concentration were assessed using the Agilent 2100 Bioanalyzer and Qubit fluorometer, and samples with RIN ≥7 were used for library preparation. Poly(A)-selected, stranded mRNA libraries were generated using BGI’s standard protocol (DNBSEQ stranded mRNA library) and sequenced as 150 bp paired-end reads on the DNBseq^TM^ platform. On average, 22 million paired-end reads.

### 2.10. RNA-seq bioinformatics analysis

Raw sequencing reads were first evaluated for quality using FastQC (Andrews S, 2010) and summarized with MultiQC (Ewels et al., 2016). Reads were aligned to the Gencode release M36 mouse reference genome (GRCm39, Ensembl release 113, primary assembly with comprehensive gene annotation) using STAR (Dobin et al., 2013). The genome index was generated with a splice junction database overhang of 149. STAR parameters included enabling two-pass mapping with an overhang of 149, setting filters for splice junctions (--alignSJoverhangMin 8, -- alignSJDBoverhangMin 1, --outFilterIntronMotifs RemoveNoncanonicalUnannotated), controlling multimapping (--outFilterMultimapNmax 20, --outMultimapperOrder Random), and applying mismatch and intron length thresholds (--outFilterMismatchNmax 999, -- outFilterMismatchNoverReadLmax 0.04, --alignIntronMin 20, --outFilterType BySJout). Gene- level quantification was performed with featureCounts (Liao et al., 2014) using exon features from the Gencode comprehensive GTF annotation, in paired-end and strand-specific mode, allowing fractional assignment of multimapping reads (-M --fraction).

Differential gene expression and functional enrichment analyses were performed within the R/Bioconductor environment v4.5.1 (R Core Team, 2021). Quality control and exploratory analyses were conducted using variance-stabilizing transformed counts (vst, blind=TRUE) from DESeq2 v1.48.1(Love et al., 2014), restricting to the 1,000 most variable genes for principal component analysis (PCA). PCA was carried out using FactoMineR v2.11 (Lê et al., 2008). Differential gene expression was assessed with DESeq2 using the Wald test to compare CD300f^−/−^ and WT samples. Fold-change estimates were corrected with the adaptive shrinkage (ashr) method (Stephens, 2016), and genes were considered differentially expressed if they had an FDR-adjusted p-value < 0.05 and an absolute log2 fold change > 0.77 after shrinkage, corresponding to a ∼70% change in expression levels. Gene annotations were retrieved using biomaRt v2.64.0 and org.Mm.eg.db v3.21.0 (Durinck et al., 2005, 2009);(Carlson M, 2025). Functional enrichment was performed with clusterProfiler v4.16.0 (Yu et al., 2012; Yu, 2024). Gene set enrichment analysis (GSEA) (Subramanian et al., 2005) was carried out on the full ranked list of genes ordered by the Wald test statistic from DESeq2. Gene sets were obtained from the Molecular Signatures Database (MSigDB) (Liberzon et al., 2011, 2015), using gene sets from the Hallmarks (MH), Reactome (M2) (Fabregat et al., 2018) and cell type signatures (M8) collections, retrieved through the msigdbr v5.1.0 package (Dolgalev I, 2025). For both ORA and GSEA, minimum and maximum gene set sizes were set to 10 and 500, respectively. Gene set variation analysis (GSVA) was performed at the individual sample level using the GSVA v2.2.0 package in R/Bioconductor (Hänzelmann et al., 2013). Normalized expression values obtained through variance-stabilizing transformation (VST, blind = FALSE) were used as input, and enrichment scores were computed after setting the gsvaParam function (kcdf = “Poisson”, minSize = 5, maxSize = 50, maxDiff = TRUE, absRanking = FALSE). Between-group comparisons of GSVA scores were performed using the Wilcoxon rank-sum test (wilcox.test(score ∼ genotype)), followed by Benjamini– Hochberg (BH) correction for multiple testing. Custom gene sets and their sources are listed in Supplementary Table 2 . The same normalized expression matrix was used to generate heatmaps with the pheatmap v1.0.13 package (Kolde R, 2025). Visualizations were generated with ggplot2 v3.5.2, gridExtra v2.3, cowplot v1.2.0, and dplyr v1.1.4 packages (Wickham et al., 2025).

### 2.11. Statistical analysis

All the results are presented as mean ± standard error of the mean (SEM). For comparison between WT and CD300f^−/−^ mice, unpaired Student’s t-test or Mann–Whitney U test was performed. For comparison between multiple groups (different time-points over the course of the cuprizone treatment), two-way ANOVA with post hoc Tukey’s or Sidak’s analysis was used. Differences were considered statistically significant when p value < 0.05, and were indicated in graphs by asterisks (*p<0.05, **p<0.01, ***p<0.001, ****p<0.0001).

## 3. RESULTS

### 3.1. Altered motor coordination after demyelination in CD300f^−/−^ mice

The CPZ model is a well-established model for studying demyelination and remyelination (Kipp et al., 2009; Ransohoff, 2012). Oral administration of the copper-chelator CPZ induces oligodendrocyte apoptosis leading to demyelination of distinct brain areas, especially the CC (Sen et al., 2020). We monitored WT and CD300f^−/−^ mice with CPZ or regular diet for 5 weeks, followed by 1 week on a regular diet, and we recorded the body weight, functional compromise, as well as neuropathologic and transcriptomic changes (Figure 1a). We observed a gradual loss of weight in animals that received CPZ diet in comparison with a regular diet, but no differences were found between animals lacking CD300f and WT mice (Figure 1b). Although, it has been established that there might be a sex-dependent effect in demyelination in the CPZ model (Taylor et al., 2009; Paton et al., 2022) and also sex-dependent effects of CD300f (Lago et al., 2020; Kaufmann et al., 2021; Evans et al., 2023), no differences between sexes were observed in our study when following body weight (Figure 1b and S1a).

To assess functional compromise, spontaneous locomotor activity was evaluated at 5 weeks of CPZ diet using the OFT. Neither CPZ treatment nor the absence of CD300f affected locomotor activity compared with animals fed with regular diet and WT mice, respectively, indicating that local demyelination does not alter the general locomotor exploratory behavior pattern of these mice (Figure 1c). Next, we evaluated motor coordination by using the grid test (Grill et al., 1997). At 5 weeks of CPZ treatment, both genotypes showed a significant increase in paw placing errors when compared with control mice (Figure 1d). Interestingly, 1 week after CPZ diet withdrawal (5+1), i.e. in the remyelination phase, WT mice tended to recover motor coordination, while CD300f^−/−^ mice displayed significantly higher number of paw-placing errors compared to WT (Figure 1d), suggesting impaired motor recovery during the remyelination phase in CD300f^−/−^ mice.

### 3.2. Absence of CD300f modifies demyelination and remyelination dynamics

Demyelination of the CC of WT and CD300f^−/−^ mice was evaluated by LFB staining after 3 and 5 weeks of CPZ treatment (Figure 2a,b). LFB staining decreased progressively in both genotypes, reaching weak myelin staining at 5 weeks of CPZ treatment. Interestingly, CD300f deficient mice showed significantly weaker myelin staining at 3 and 5 weeks of CPZ diet than WT mice (Figure 2a,b), suggesting a higher degree of demyelination. After 1 week of CPZ withdrawal, LFB staining began to recover in WT mice, but this recovery was significantly lower in CD300f^−/−^ mice, suggesting a delay in the remyelination process in both males (Figure 2a,b) and females (Supplementary Figure S1b).

**Figure 2:**
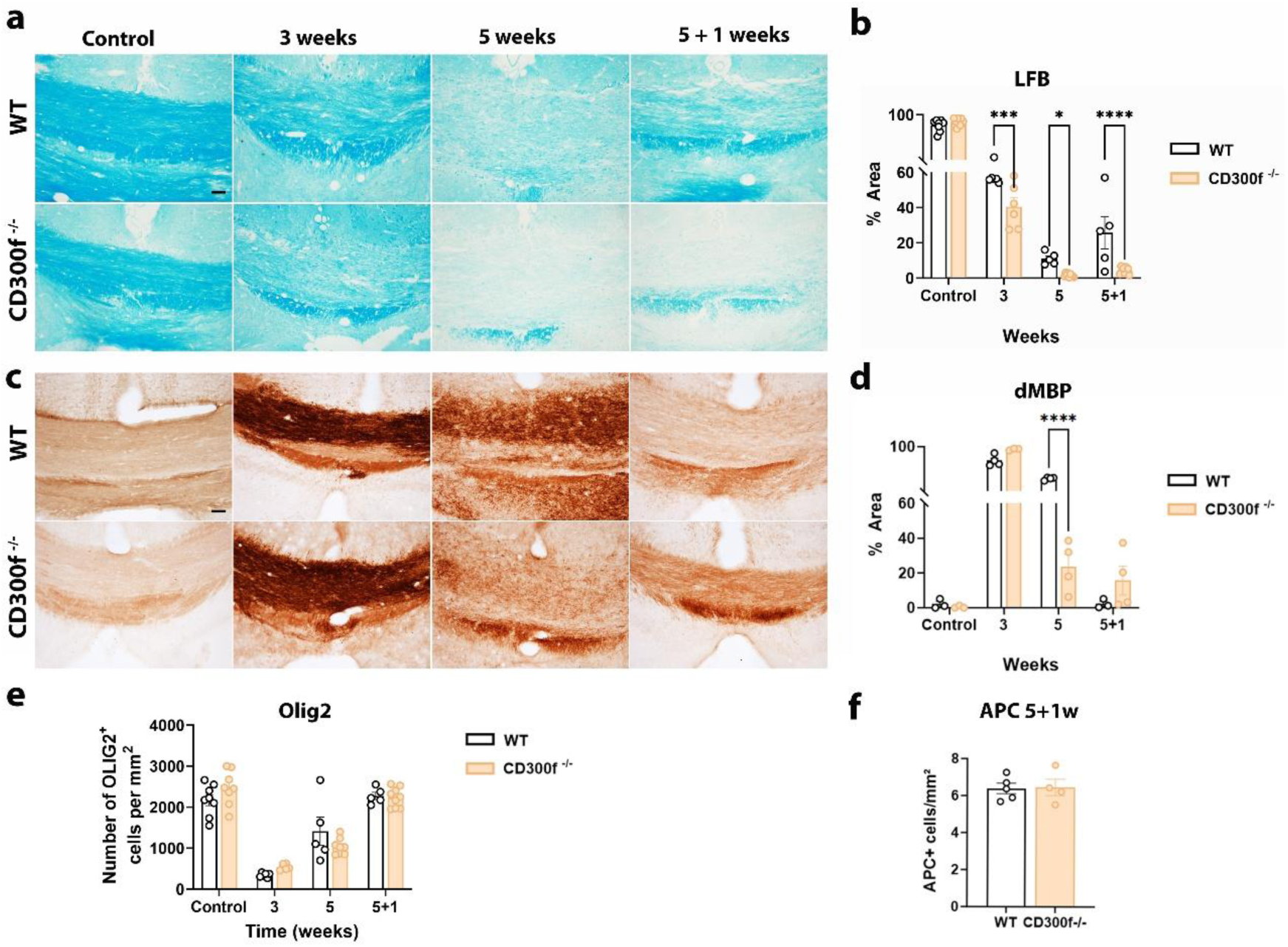
CD300f^−/−^ mice show increased demyelination and impaired remyelination. **(a-b)** Representative images of Luxol Fast Blue (LFB) staining **(a)** in the central corpus callosum (CC) and the corresponding quantification of the LFB-stained area (% Area) **(b)** in control, 3- and 5-week CPZ-treated WT and CD300f^−/−^ mice, and after 1-week CPZ treatment withdrawal (5+1) in both WT and CD300f^−/−^ mice. **(c-d)** Representative images of dMBP immunostaining at the central CC **(c)** and the corresponding quantification of the dMBP immunolabeled area (% Area) **(d)** in control, 3- and 5-week CPZ-treated WT and CD300f^−/−^ mice, and after 1-week CPZ treatment withdrawal (5+1) in both WT and CD300f^−/−^ mice. **(e)** Quantification of the oligodendroglial population, stained as Olig2-positive cells (Olig2+/mm^2^) in control, 3- and 5-week CPZ-treated WT and CD300f^−/−^ mice, and in mice after 1 week of CPZ withdrawal (5+1). (**f)** Quantification of mature oligodendrocytes, stained as APC-positive cells (APC+ cells/mm^2^), after 1 week of CPZ withdrawal (5+1), during remyelination. Data are presented as mean ± SEM. Each dot represents an animal. Statistical analysis: Two-way ANOVA plus Tukey’s post-hoc test and two-tailed Student’s t- test (APC 5+1); *p≤0.05, ***p≤0.001, ****p≤0.0001. Scale bar (a, c): 50 µm.

To further assess the extent of myelin debris or aberrant myelin accumulation in the CC, we analyzed degraded myelin basic protein (dMBP) using an antibody that binds MBP epitopes exposed only after myelin degradation (Figure 2c,d). After 3 weeks of CPZ-treatment maximum and similar levels of dMBP staining were observed in both genotypes reflecting MBP degradation. However, at 5 weeks of CPZ treatment, dMBP levels were slightly reduced in WT mice, while CD300f^−/−^ mice showed a stronger, significant reduction. One week after CPZ withdrawal, dMBP decreased significantly in WT mice returning to control levels, while similar levels remained in CD300f^−/−^ mice (Figure 2c,d).

CPZ treatment specifically causes the death of mature oligodendrocytes, and therefore, relevant changes are observed in the dynamics of the whole oligodendrocyte population in this model (Petković et al., 2016). After 3 weeks of CPZ treatment, the overall OLIG2^+^ population was dramatically reduced in both genotypes when compared with control mice (Figure 2e). The OLIG2^+^ cell population began to slightly recover after 5 weeks of CPZ treatment, reaching normal values during the remyelinating phase (5+1 weeks) in both genotypes and sexes (Figure 2e; Supplementary Figure S2c). Moreover, no differences in the number of mature oligodendrocytes stained with the APC marker were observed between genotypes during the remyelination phase (Figure 2f).

Taken together, these results suggest that CD300f deficiency enhances demyelination and lipid disorganization during the demyelination phase (Figure 2a-d), potentially contributing to the motor impairment observed during the remyelination phase (Figure 1d). Moreover, during remyelination, CD300f deficiency may hamper the reemergence of structured lipid myelin components, without affecting either MBP degradation levels or the general oligodendrocyte dynamics.

### 3.3. Absence of CD300f delays microglial response to demyelination

Microglial and astrocytic changes were analysed through the immunohistochemical detection of Iba-1 and GFAP, respectively. It has been demonstrated that the CPZ diet induces demyelination without recruitment of peripheral monocytes (Mildner et al., 2007); therefore, Iba1 staining reflects microglial dynamics exclusively. CPZ treatment progressively increased microglial reactivity, peaking at 5 weeks. Interestingly, at this time-point, CD300f^−/−^ mice exhibited significantly lower microglial Iba1-positive staining area (Figure 3a,b) than WT mice. Analysis of PU.1+ cells revealed a trend toward fewer microglia in CD300f^−/−^ mice compared with WT at 5 weeks, consistent with the reduction observed in the Iba1-positive area, although the difference did not reach statistical significance (Figure 3c,d). After one week of remyelination, microglial reactivity decreased similarly in both genotypes, although it did not return to control levels (Figure 3a,b).

**Figure 3:**
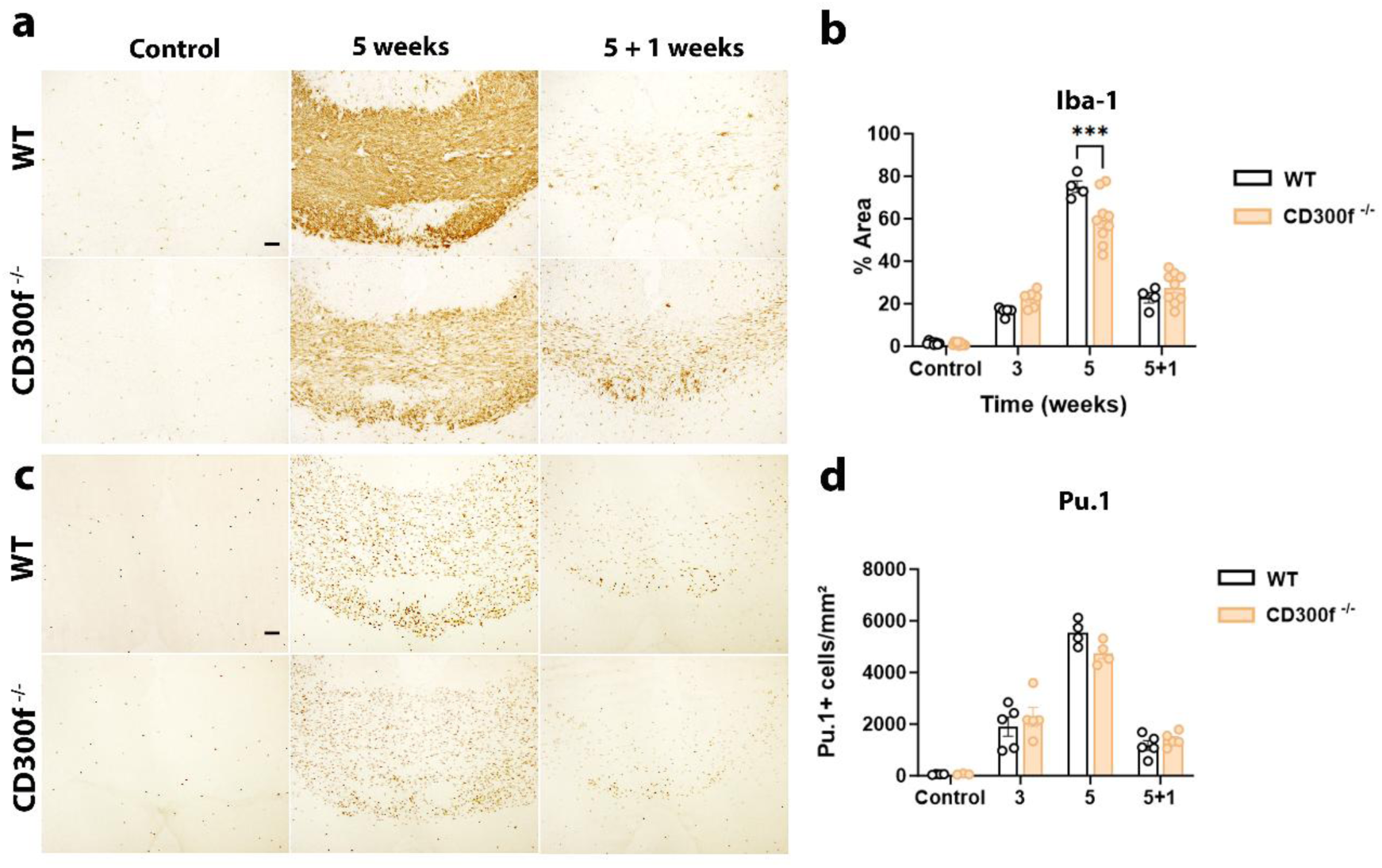
CD300f^−/−^ exhibits reduced microglial reactivity during demyelination. **(a-b)** Representative images of Iba-1 immunostaining **(a)** in the central corpus callosum (CC) of control, 5-week CPZ-treated WT and CD300f^−/−^ mice, and after 1 week of CPZ withdrawal (5+1). Quantification of the Iba-1 stained area (% Area) of both genotypes after CPZ treatment is shown in **(b)**. **(c)** Representative images of Pu.1 immunostaining for detection of microglia cell number in the central CC of control, 5-week CPZ-treated WT and CD300f^−/−^ mice, and after 1 week of CPZ withdrawal (5+1). **(d)** Quantification of the Pu.1-positive cell density in the central CC of both genotypes after CPZ treatment. Data are presented as mean ± SEM. Each dot represents an animal. Statistical analysis: Two-way ANOVA plus Tukey’s post-hoc test: ***p≤ 0.001. Scale bar (a, c): 50 µm.

After CPZ diet, GFAP expression progressively increased up to 5 weeks of treatment, remaining at high levels after 1 week of remyelination, without relevant differences between genotypes (Supplementary Figure 2).

### 3.4 Loss of CD300f amplifies microglial and inflammatory transcriptional programs during demyelination

Given CD300f expression in central nervous system microglia, we used bulk RNA-seq to investigate how its deletion affects the CC transcriptional response to cuprizone-induced demyelination, with particular attention to microglia-associated changes. CD300f^−/−^ and WT mice were fed a CPZ diet for 5 weeks, the peak of demyelination, and the CC was isolated and processed for bulk transcriptomics (Supplementary Table 3). CD300f^−/−^ and WT mice were fed with CPZ diet for 5 weeks, reaching the peak of demyelination, and the CC was isolated and processed for bulk RNA-seq. PCA revealed a clear separation between CD300f^−/−^ and WT samples along PC1, which explained 64.6% of the total variance, while PC2 accounted for an additional 11.5%, (Figure 4a). Differential expression analysis identified 207 genes meeting significance thresholds (FDR-adjusted p-value < 0.05 and |log2FC| > 0.77 after ashr shrinkage), with changes in both directions (Figure 4b and Supplementary Table 4).

**Figure 4.**
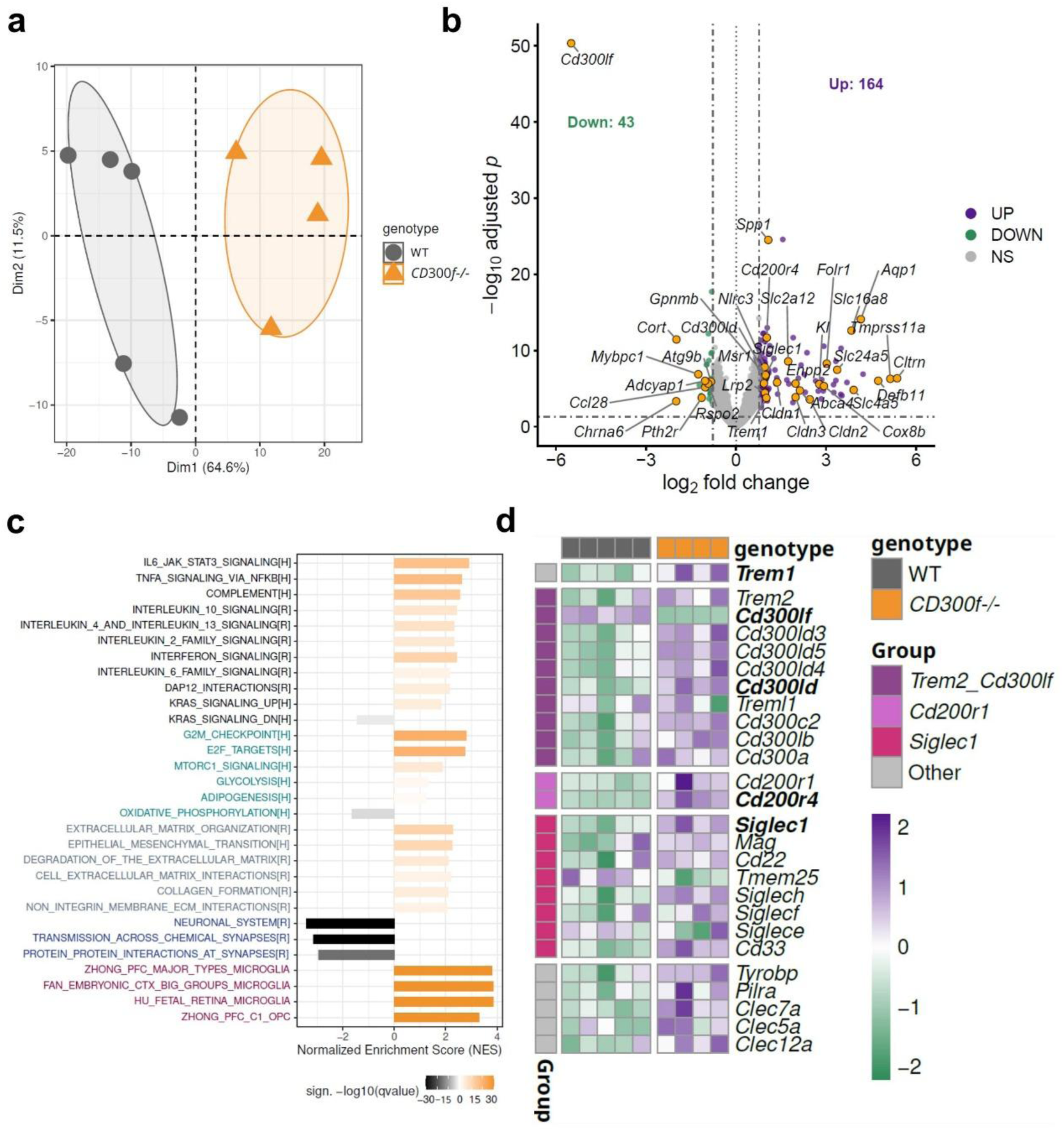
Transcriptomic profiling reveals myeloid and matrix-remodeling signatures in the CD300f^−/−^ corpus callosum following a 5-week cuprizone diet. **(a)** Principal component analysis (PCA) of variance-stabilizing transformed (vst, blind = TRUE) expression values for the 1,000 most variable genes, showing separation of CD300f^−/−^ and WT samples. **(b)** Volcano plot of differential expression results (DESeq2 Wald test; adaptive shrinkage of log₂ fold changes with ashr). Genes with FDR-adjusted p-value < 0.05 and absolute ashr-shrunken log2 fold change > 0.77 are highlighted in purple (upregulated) and green (downregulated). *Cd300lf*, the targeted gene, is the most strongly downregulated transcript. *Cd300lf* and additional genes of interest are highlighted in orange. **(c)** Normalized enrichment scores (NES) obtained from gene set enrichment analysis (GSEA) using the MSigDB Hallmark [MH], Reactome [MR], and cell-type signature collections [M8]. Selected significantly enriched gene sets are shown. Hallmark and Reactome gene set labels are colored according to functional category: black, inflammatory; turquoise, metabolic and proliferative; gray, extracellular matrix (ECM)-related; and blue, neuronal or synaptic. Cell-type signatures are labeled in purple. Bar length and direction represent the magnitude and sign of the NES, respectively, whereas bar color represents the signed −log10(q-value), with the sign determined by the direction of enrichment. **(d)** Heatmap showing the expression of selected immune receptor-related genes, including members of the Cd300, Trem, Cd200r, and Siglec families and the signaling adaptor *Tyrobp*. CD300f^−/−^ samples generally showed higher expression of these genes than WT samples. Variance-stabilizing transformation (vst; blind = FALSE) of expression values was standardized by gene and displayed as row Z-scores. Rows represent genes and columns represent individual samples. Genes meeting both differential expression criteria (FDR-adjusted p-value < 0.05 and |ashr-shrunken log2 fold change| > 0.77) are shown in bold: *Cd300lf*, *Cd300ld*, *Cd200r4*, *Trem1*, and *Siglec1*. Among the remaining genes, *Tmem25* was significantly downregulated, whereas *Trem2*, *Cd300lb*, *Cd300lc2*, *Cd300ld3*, *Cd300ld4*, *Cd300ld5*, *Cd200r1*, *Siglech*, *Cd33*, *Tyrobp*, *Pilra*, *Clec5a*, *Clec7a* and *Clec12a* were significantly upregulated (FDR-adjusted p-value < 0.05), but did not exceed the absolute log2 fold-change threshold of 0.77. *Cd300lf* is included as a reference.

Gene set enrichment analysis (GSEA) across the full ranked gene list revealed a coordinated upregulation of immune-related programs in CD300f^−/−^ mice, encompassing interferon responses, pro-inflammatory cytokine and chemokine signaling (TNFα/NF-κB, IL6/JAK/STAT3), complement activation, and anti-inflammatory cascades (IL-10, IL-4/IL-13), together with enrichment of the DAP12 signaling pathway, a central node of myeloid activation (Figure 4c and Supplementary Table 5) (Hickman et al., 2013). The increased inflammatory response was accompanied by downregulation of the oxidative phosphorylation metabolic program, whereas glycolysis, mTORC1 signaling, and adipogenesis showed milder upregulation (Figure 4c), collectively suggesting a metabolic shift accompanying myeloid activation. The pyroptosis *Casp1/Nlrp3/Gsdmd* and apoptotic *Casp8/Casp3* mRNAs were upregulated, suggesting that some cells may be undergoing cell death such as OPC/NFO or microglia.

Signatures related to neuronal and synaptic function were enriched among the downregulated genes (Figure 4c and Supplementary Table 5). This may reflect decreased mRNA transport along axons due to increased axonal loss, or changes in gene expression attributable to gray-matter contamination in the CC samples. Finally, several immune receptors were upregulated in CD300f^−/−^ mice, including members of the CD300, Trem, CD200R, and sialic acid–binding receptor families, together with the signaling adaptor DAP12 encoded by *Tyrobp* gene. This expression pattern is consistent with an enhanced representation of myeloid receptor–adaptor modules in the absence of CD300f (Figure 4d).

Consistent with this immune-skewed profile, GSEA using MSigDB cell-type signatures revealed significant positive enrichment of gene sets associated predominantly with microglia and other myeloid populations in the knockout condition, supporting a prominent contribution of microglia-associated transcriptional changes to the bulk RNA-seq response (Figure 4c).

Gene Set Variation Analysis (GSVA) of corpus callosum samples revealed significantly higher scores in CD300f^−/−^ mice for multiple microglial signatures, including both homeostatic and disease-associated microglia (DAM) programs (Figure 5a). Complementary GSEA using the same gene sets yielded concordant enrichment patterns (Supplementary Figure 3a, and Supplementary Table 6). This pattern indicates the concurrent representation of distinct microglial transcriptional states in the CD300f-depleted condition. CD300f^−/−^ samples also exhibited significantly higher scores for several demyelination-associated microglial signatures, including those identified in the cuprizone model by Hou et al. (2023) (Hou et al., 2023). Moreover, a distinct population of foamy GPNMB+ microglia/macrophages has been associated with lesion expansion in secondary progressive MS (van der Vliet et al., 2026). The corresponding transcriptional signature showed significantly higher GSVA scores in CD300f^−/−^ samples, indicating greater representation of this expression program in the knockout condition. GSVA revealed higher scores in CD300f^−/−^ samples for gene sets associated with lipid recognition and uptake, efferocytosis, lysosomal and phagosomal function, and cholesterol metabolism (Figure 5b, Supplementary Figure 3b and Supplementary Table 6). Consistent with these pathway-level results, differential expression analysis identified increased expression of several genes involved in these processes (Supplementary Table 4). These included lipid-sensing, scavenger, and uptake receptors such as *Cd36*, *Apobr*, *Msr1*, *Lrp1*, *and Trem2*, as well as the efferocytosis receptors *Mertk* and *Axl* and their respective ligands *Pros1* and *Gas6*. Genes involved in lysosomal lipid degradation were also upregulated in CD300f^−/−^ mice, including *Lipa*, *Gba1*, *Galc*, *Smpd1*, and *Asah1*, together with the lysosomal cholesterol transporter *Npc2* and several cathepsins, including *Ctsb*, *Ctsd*, *Ctse*, and *Ctsz* (Figure 5b; Supplementary Table 4). In contrast, the cholesterol biosynthesis program was reduced, consistent with the decreased expression of *Hmgcr* and *Hmgcs1*, whereas no comparable change was detected in fatty acid synthesis. CD300f^−/−^ samples also showed increased expression of genes involved in lipid uptake, processing, transport, and storage, including *Apoc1*, *Apoe*, *Ch25h*, *Fabp5*, *Lpl*, and *Soat1*. Moreover, the cholesterol efflux transporters *Abca1* and *Abcg1* were upregulated, along with genes involved in intracellular cholesterol trafficking and esterification, including *Npc2* and *Soat1*. To validate these findings, we visualized the accumulation of digested myelin in lipid droplets using Oil Red O (ORO) staining of triglycerides (TAGs) and cholesteryl esters (Mehlem et al., 2013). After 5 weeks of CPZ treatment, lipid droplets appeared in the CC. Interestingly, CD300f^−/−^ mice showed a marked increase in the ORO-positive area, denoting greater lipid droplet accumulation in the absence of CD300f (Figure 5d-e), particularly within microglial cells, as revealed by BODIPY co-staining (Figure 5f). Together with the observed changes in gene expression, these results may reflect an impaired ability of CD300f^−/−^ microglia to cope with lipid accumulation during demyelination.

**Figure 5.**
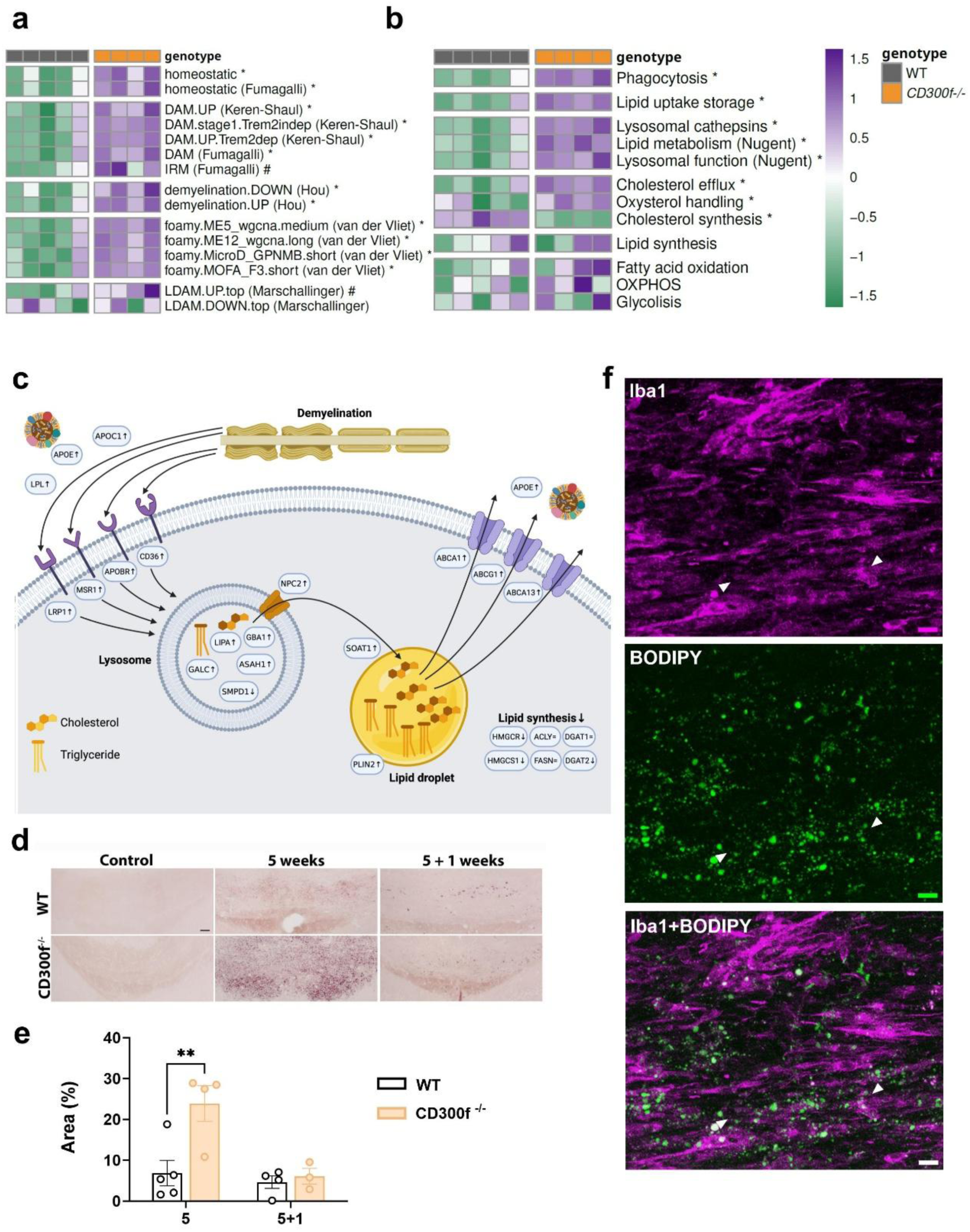
Altered microglial transcriptomic programs and lipid handling during demyelination in CD300f^−/−^ mice. **(a-b)** Gene set variation analysis (GSVA) of bulk RNA-seq from CC of CD300f^−/−^ (n = 4) and wild-type (n = 5) mice at the 5-week demyelination stage. **(a)** Enrichment scores for curated microglial state signatures, obtained from (Keren-Shaul et al., 2017; Marschallinger et al., 2020; Hou et al., 2023; Fumagalli et al., 2025; van der Vliet et al., 2026). **(b)** Enrichment scores for metabolism-related gene sets, including lipid handling-related programs and gene sets from Nugent et al., 2020. Each column is a sample and each row a gene set; color indicates the row-scaled (z-scored) GSVA enrichment score (see color key). Differences between CD300f⁻ᐟ⁻ and wild-type were assessed per gene set by Wilcoxon rank-sum test with Benjamini–Hochberg correction: *adjusted p-value < 0.05; #adjusted p-value < 0.10. Gene sets used in a) and b) are defined in Supplementary Table 2. **(c)** Representative scheme showing alterations in the expression (arrows indicate direction) of key lipid receptors, lysosomal enzymes, lipid droplet components or cholesterol export machinery in CD300f^−/−^ mice. **(d)** Representative images of the staining of CC at 5 weeks of CPZ diet and 1 week after CPZ withdrawal with ORO for neutral lipids and quantification showed increased lipid droplets in CD300f^−/−^ mice **(e)**. Data are presented as mean ± SEM. Each dot represents an animal. Statistical analysis: Two-way ANOVA followed by Sidak post-hoc test; **p ≤ 0.01. Scale bar (a, c): 50 µm. **(f)** Representative confocal images of double immunostaining for Iba1 (magenta) and BODIPY (green) in CD300f^−/−^ mice after 5 weeks of CPZ diet. BODIPY-positive lipid droplets were predominantly associated with Iba1-positive microglia, although some lipid droplets were detected outside microglial cells (arrowheads). Scale bar (f): 10 µm.

## 4. DISCUSSION

In the present study, we investigated the role of the immune receptor CD300f in demyelination and remyelination using the CPZ model. CD300f deficiency exacerbated demyelination after 5 weeks of CPZ treatment, as evidenced by reduced LFB staining, lower levels of degraded MBP (dMBP), and reduced microglial density. During the remyelination phase, CD300f-deficient mice displayed reduced myelin recovery accompanied by impaired motor coordination. Transcriptomic analysis at the peak of demyelination revealed enrichment of homeostatic and DAM microglial signatures together with increased expression of genes involved in phagocytosis, inflammation, lysosomal activity, and lipid metabolism. These alterations were accompanied by a significant upregulation of foamy microglial signatures previously associated with demyelination (van der Vliet et al., 2026), accompanied by marked lipid droplet accumulation. Collectively, these findings indicate that CD300f promotes effective microglial responses to myelin damage and contributes to efficient tissue repair following demyelination.

It is well established that CPZ induces apoptosis of mature oligodendrocytes, resulting in CNS demyelination, particularly in the CC, accompanied by astrocyte and microglial activation (Kipp et al., 2009; Skripuletz et al., 2013; Sen et al., 2022). In this context, the more severe demyelination after 5 weeks of CPZ treatment that extends during the remyelination phase suggests that demyelination is not only more pronounced but also remains actively ongoing in CD300f-deficient mice at the peak of disease. Increased myelin degeneration may reflect enhanced detection and/or clearance of myelin debris, processes that are essential for limiting secondary tissue damage and enabling subsequent remyelination. However, despite this accelerating clearance and tissue repair, CD300f deficiency ultimately impaired both remyelination and functional recovery. CD300f has been proposed to function as a DAMP receptor involved in sensing tissue injury *in vivo*, although the underlying mechanisms remain incompletely understood (Negro-demontel et al., 2024). Several lipid species, including sphingomyelin and phosphatidylserine, act as CD300f ligands and may contribute to microglial recognition of tissue damage and coordination of the subsequent cellular response. In addition to myelin detection and engulfment, successful adaptation to chronic demyelination requires efficient metabolic processing of internalized material. Previous studies have shown that CD300f-deficient microglia exhibit reduced metabolic fitness under inflammatory conditions, characterized by decreased expression of glucose transporters, glycolytic enzymes, lipid transporters, and β-oxidation pathways (Lago et al., 2020). Likewise, CD300f-deficient macrophages and microglia display a reduced capacity to metabolize engulfed apoptotic material (Negro-demontel et al., 2024). Consistent with this notion, our transcriptomic data suggest reduced oxidative phosphorylation and increased glycolytic pathways in the CC of CD300f-deficient mice. However, because these analyses were performed on bulk tissue, the cellular origin of these changes cannot be determined. Future single-cell and spatial transcriptomic studies will be necessary to establish how CD300f deficiency affects distinct microglial populations and other CNS cell types during demyelination.

Despite the reduced microglial density and prominent lipid droplet accumulation, the CC of CD300f^−/−^ mice displayed increased expression of genes involved in lysosomal activity (*Cst7, Cstb, Ctsd, Ctse, Ctsl*), lipid metabolism (*Apoc1, Apoe, Lipa, Lpl*), cholesterol handling (*Abca1, Abcg1, Soat1*), phagocytosis (*Trem2, Mertk, Csf1r, Spp1, CD86, CD68*), and inflammatory responses, together with markers associated with both disease-associated and homeostatic microglial states. This transcriptional profile suggests that the remaining microglia are exposed to persistent myelin damage and attempt to compensate through enhanced degradative and lipid-processing programs. Nevertheless, the concomitant accumulation of lipid droplets indicates that these responses are insufficient to restore lipid homeostasis, resulting in lipid overload and metabolic stress that may further compromise microglial fitness and survival. Importantly, as transcriptomic analyses were performed on bulk CC tissue, the observed gene expression changes cannot be attributed specifically to microglia and may reflect a combination of cell-intrinsic changes in microglia and other glial populations. Although we cannot rule lipid droplets accumulation also in astrocytes, when integrated with the histological and functional data, the transcriptomic signature is consistent with these processes occurring mainly in microglial cells. This lipid droplet accumulation is increasingly recognized as a hallmark of altered lipid processing in microglia, particularly under conditions of excessive myelin degradation (Marschallinger et al., 2020). Cholesterol accumulation has been described in phagocytes from aged mice with EAE following excessive uptake of myelin debris (Cantuti-Castelvetri et al., 2018). More recently, a population of foamy GPNMB+ microglia/macrophages was identified in progressive MS lesions and associated with lesion expansion, accumulation of cholesterol esters and triglycerides, lysosomal stress, and enhanced phagocytic activity (van der Vliet et al., 2026). In agreement with these observations, GSVA and complementary GSEA analysis of CD300f-deficient mice revealed enrichment of both the GPNMB+ and DAM-like microglial states defined in that study. Notably, integrative lipidomic, transcriptomic, and proteomic analyses linked the GPNMB+ population to disease progression and identified CD300f among the major transcriptomic contributors associated with this pathogenic signature (van der Vliet et al., 2026).These findings strengthen the association between CD300f biology, foamy microglial states, and chronic demyelinating pathology. Further mechanistic studies are required to determine whether these lipid-laden microglia actively drive disease progression and to define the precise role of CD300f across different microglial and macrophage states.

Other lipid droplet-associated microglial states have been described during aging, such as LDAMs (Marschallinger et al., 2020). However, our data do not support a similar mechanism. In LDAM, the rate-limiting enzyme in fatty acid and cholesterol synthesis *Acly* is induced, leading to accumulation of glycerolipids such as triacylglycerol but not cholesterol. We do not observe induction in *Acly* or *Fasn* (fatty acid synthase) and detected a reduction in *Hmgcr* (HMG-CoA reductase), the rate limiting enzyme of cholesterol synthesis. In contrast, we observed upregulation of genes involved in lipid uptake, storage, and cholesterol efflux, including *Soat1, Plin2, Abca1, Abca13, and Abcg1*. Together, these observations suggest that the lipid accumulation observed in CD300f^−/−^ mice more closely resembles the cholesterol-rich foamy phenotype described in EAE and MS than the aging-associated LDAM phenotype. Importantly, despite activation of cholesterol efflux pathways, lipid droplets continued to accumulate, suggesting that lipid storage and handling mechanisms remain insufficient to compensate for the increased burden of myelin-derived lipids.

The role of CD300f in phagocytosis and lipid metabolism remains poorly understood. Although CD300f is not required for non-receptor-mediated phagocytosis of inert particles by microglia (Lago et al., 2020), it has been implicated in apoptotic cell recognition and clearance (Tian et al., 2014; Negro-demontel et al., 2024) as well as LDL uptake (Lago et al., 2020). Our results do not demonstrate a direct role for CD300f in myelin phagocytosis, but they support an important function for this receptor in coordinating the metabolic adaptations required for efficient handling of myelin-derived lipids during demyelination. Interestingly, the functionally relevant R218Q CD300f variant has previously emerged as a putative susceptibility locus for multiple sclerosis, although it did not reach genome-wide significance (Ban et al., 2010). Moreover, CD300f surface expression was significantly reduced on classical and intermediate monocytes in MS patients compared with healthy individuals (Martínez-Barriocanal et al., 2017). Together, these findings provide functional evidence supporting a role for CD300f in demyelinating disease and warrant further investigation of the underlying molecular mechanisms.

An interesting observation was the reduced microglia density despite the increased burden of myelin damage. Efficient clearance and processing of myelin debris rely on the recruitment, survival, and metabolic adaptation of microglia within demyelinated lesions. The reduced microglial density observed in CD300f-deficient mice may therefore reflect impaired cellular metabolic fitness and increased susceptibility to cell death. Supporting this possibility, transcriptomic analyses revealed increased expression of *Casp8* and *Casp3*, suggesting activation of apoptotic pathways, as well as NLRP3, *Casp1*, and *Grzmb*, which may indicate pyroptotic signaling. These observations align with recent reports showing enhanced pyroptosis in CD300f-deficient microglia under inflammatory conditions (Chen et al., 2026), and with previous findings of reduced microglial numbers and increased lipid droplet accumulation in aged CD300f^−/−^ mice (Evans et al., 2023). A similar pattern of fewer Iba1+ cells has been previously observed in mice lacking TREM2 during aging or at 4 weeks of CPZ treatment (Poliani et al., 2015). Because we observed no changes in astrocyte and Olig2 or APC oligodendroglia cell densities in CD300f-deficient mice, these data may reflect microglial cell death. However, direct validation of these cell death pathways was beyond the scope of the present work and will require future investigation.

Importantly, demyelination in the CPZ model is primarily driven by oligodendrocyte toxicity rather than adaptive immune mechanisms (Kipp et al., 2009). Therefore, the exacerbated demyelination observed in CD300f-deficient mice is unlikely to result from increased autoimmune inflammatory destruction. Instead, reduced microglial abundance and altered metabolic adaptation may impair the efficient clearance and processing of myelin debris, prolonging tissue damage and ultimately compromising remyelination. The impaired motor coordination observed during the remyelination phase further supports the notion that CD300f contributes not only to structural repair but also to functional recovery following myelin injury. Similar deficits in functional recovery have been reported in other models of CNS injury involving CD300f deficiency such as traumatic brain injury (Lu et al., 2024; Negro-demontel et al., 2024). Together, our findings support a model in which CD300f promotes microglial adaptation to myelin damage, facilitating lipid homeostasis, tissue repair, and successful remyelination.

## 5. CONCLUSIONS

In summary, our data identify CD300f as a critical regulator of microglial adaptation to myelin-derived lipid stress during demyelination. Structurally and functionally related to TREM2 yet mechanistically distinct, CD300f appears to be required to couple lipid sensing with efficient phagocytosis, metabolic reprogramming, and resolution of inflammation. In its absence, microglia mount a robust but maladaptive transcriptional response like the one observed in a subset of microglia in MS progressing lesions characterized by enhanced lipid uptake, lysosomal processing and lipid droplet storage that ultimately fails to restore CNS homeostasis. These findings underscore the importance of lipid-sensing immune receptors in shaping microglial function and highlight CD300f as a potential target for modulating microglial responses in demyelinating diseases.

## Acknowledgments

The authors thank Miguel Martil for technical assistance with the histological studies.

## Author contributions

GM, NL, and HP conceived the study. GM, NR, EF, PSM and NL performed the experiments. GM, NR, BA, RLV, HP and NL analysed and discussed results. GM, NR, HP and NL drafted the original manuscript and PSM, NR, RLV, BC, BG, BA, HP, GM and NL contributed to the edition of the final version. All authors read and approved the final manuscript.

## Data availability statement

The raw RNA-sequencing data generated in this study have been deposited in the NCBI Sequence Read Archive (SRA) under BioProject accession number PRJNA1520429. All non-transcriptomic data generated in this study are available from the corresponding author upon reasonable request. No specific code was generated for this work.

---

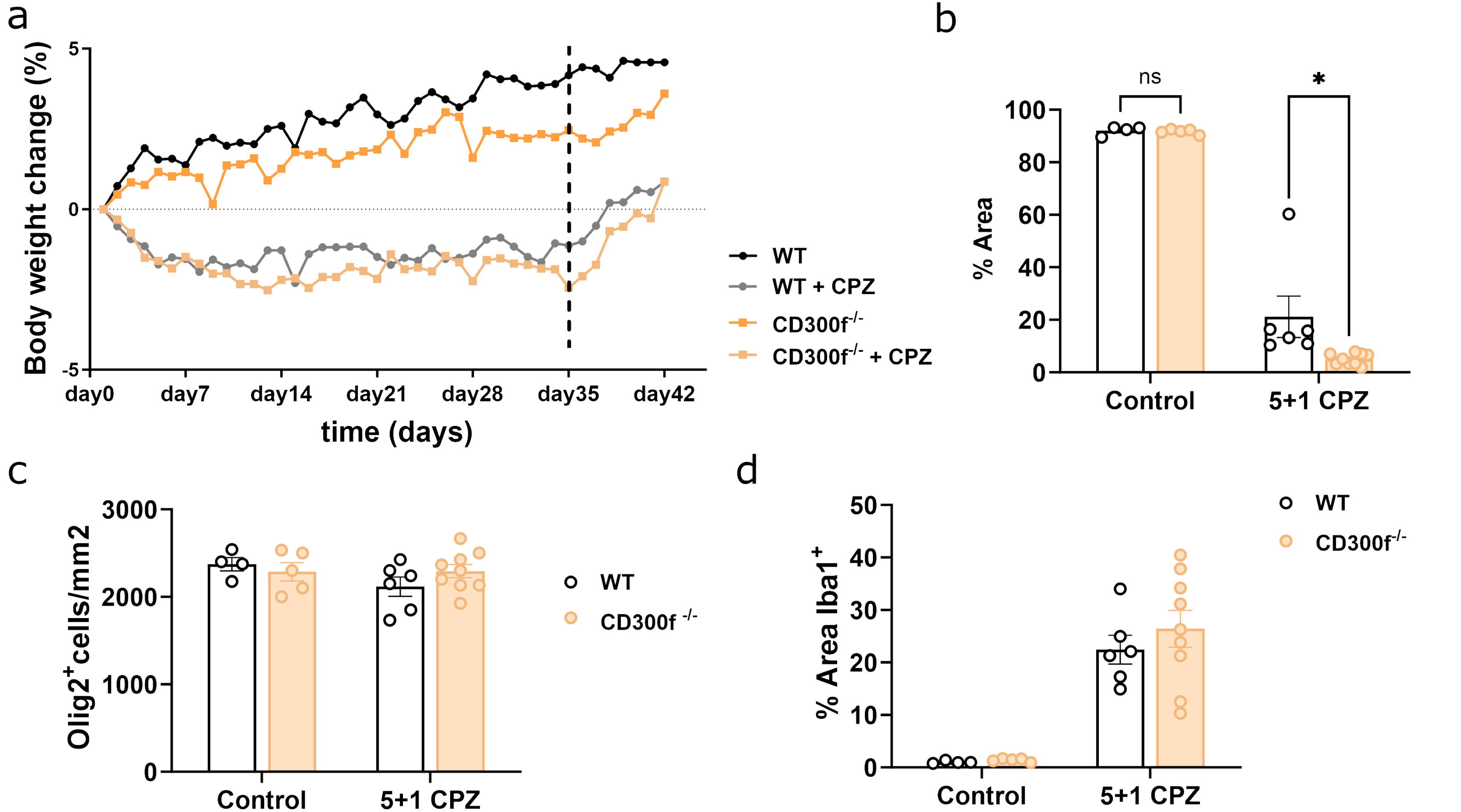

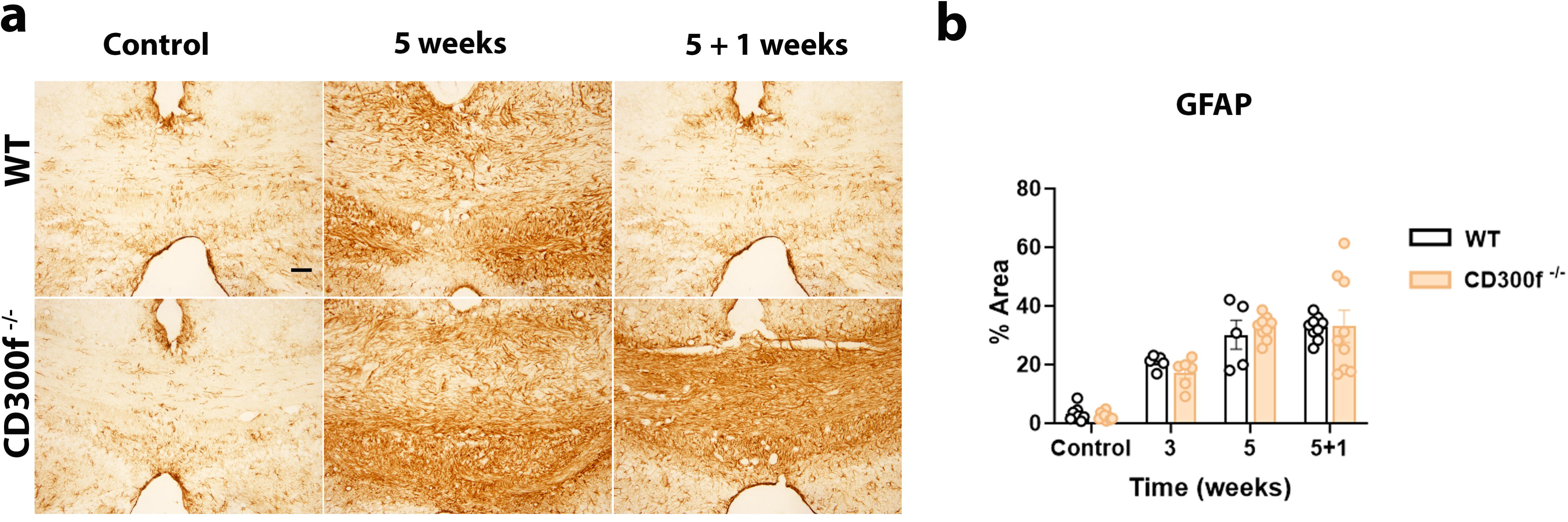

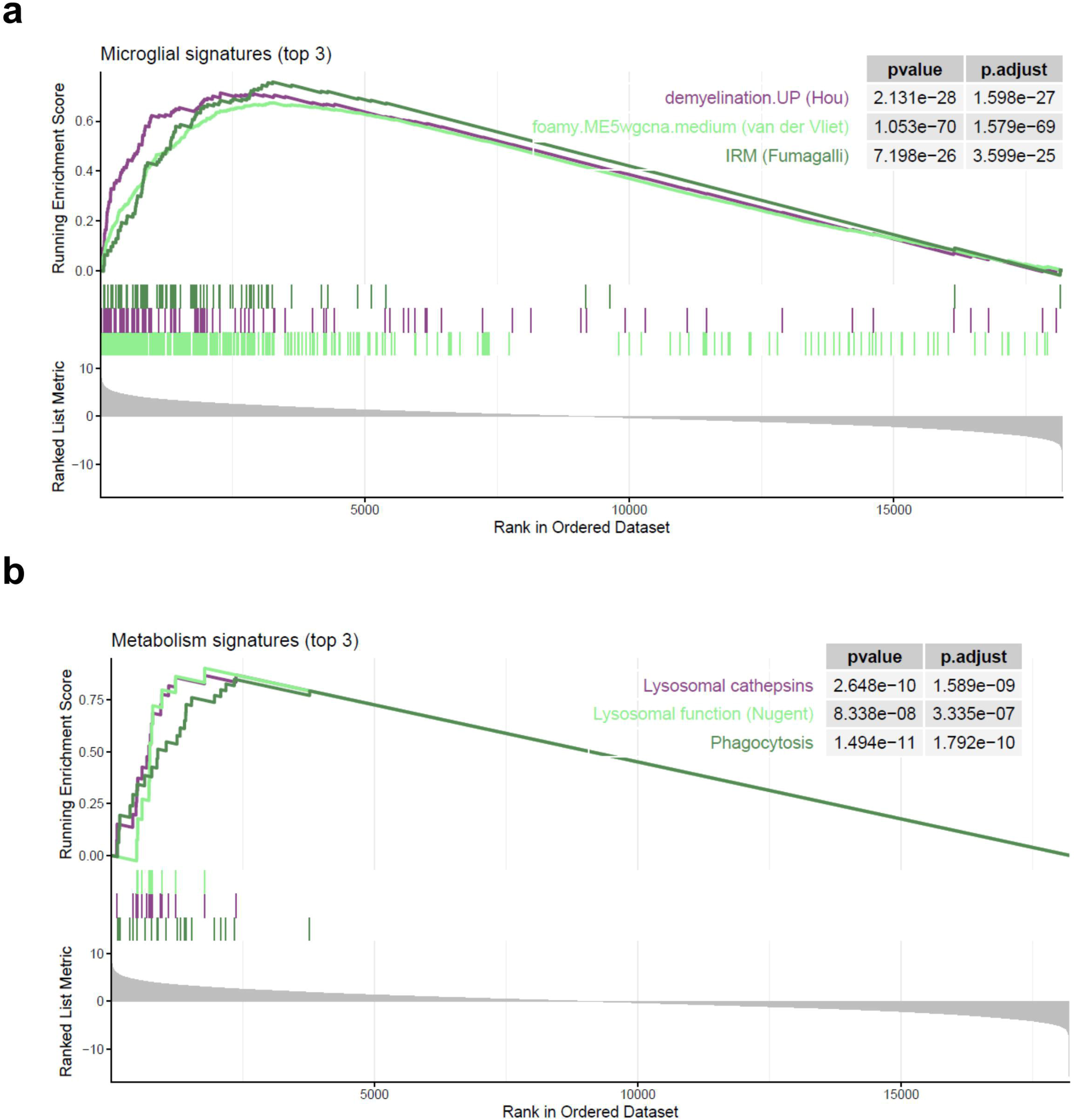

